# An integrated metagenomics pipeline for strain profiling reveals novel patterns of transmission and global biogeography of bacteria

**DOI:** 10.1101/031757

**Authors:** Stephen Nayfach, Beltran Rodriguez-Mueller, Nandita Garud, Katherine S. Pollard

## Abstract

We present the *Metagenomic Intra-species Diversity Analysis System (MIDAS)*, which is an integrated computational pipeline for quantifying bacterial species abundance and strain-level genomic variation, including gene content and single nucleotide polymorphisms, from shotgun metagenomes. Our method leverages a database of >30,000 bacterial reference genomes which we clustered into species groups. These cover the majority of abundant species in the human microbiome but only a small proportion of microbes in other environments, including soil and seawater. We applied *MIDAS* to stool metagenomes from 98 Swedish mothers and their infants over one year and used rare single nucleotide variants to reveal extensive vertical transmission of strains at birth but colonization with strains unlikely to derive from the mother at later time points. This pattern was missed with species-level analysis, because the infant gut microbiome composition converges towards that of an adult over time. We also applied *MIDAS* to 198 globally distributed marine metagenomes and used gene content to show that many prevalent bacterial species have population structure that correlates with geographic location. Strain-level genetic variants present in metagenomes clearly reveal extensive structure and dynamics that are obscured when data is analyzed at a higher taxonomic resolution.

## Introduction

Microbial communities play a myriad of important roles in the different environments that they inhabit. These communities are typically comprised of various distinct species that each exists as a complex and heterogeneous population of cells with differences in gene content (Greenblum et al. 2015; Zhu et al. 2015) and single nucleotide polymorphisms (SNPs) (Schloissnig et al. 2013; Kashtan et al. 2014; Lieberman et al. 2014). Several recent studies used within-species differences in gene content and SNPs as a window into the ongoing evolutionary history of microbes on earth. For example, this approach has revealed genomic events that lead to ecologically distinct species (Shapiro et al. 2012), uncovered the presence of ancient subpopulations of ecologically differentiated marine bacteria (Kashtan et al. 2014), and highlighted extensive intra-species recombination in pathogens (Snitkin et al. 2011) and free-living bacteria (Rosen et al. 2015). Additionally, an understanding of strain-level variation is critical for studying the interaction of microbes with humans and for understanding microbial pathogenicity. Differences at the nucleotide level can lead to within-host adaptation of pathogens (Lieberman et al. 2014), and differences in gene content can confer drug resistance, convert a commensal bacterium into a pathogen (Snitkin et al. 2011), or lead to outbreaks of highly virulent strains (Rasko et al. 2011).

Metagenomic shotgun sequencing has the potential to shed light onto strain-level heterogeneity among bacterial genomes within and between microbial communities, yielding a genomic resolution not achievable by sequencing the 16S ribosomal RNA gene alone (Sunagawa et al. 2013) and circumventing the need for culture-based approaches. However, limitations of existing computational methods and reference databases have prevented most researchers from obtaining this level of resolution from metagenomic data. Assembly-free methods that map reads to reference genomes in order to estimate the relative abundance of known strains (Francis et al. 2013; Tu et al. 2014) are effective for well-characterized pathogens like *E. coli* that have thousands of sequenced genomes. However, such methods cannot detect strain-level variation for the vast majority of known species that currently have only a single sequenced representative. Other assembly-free approaches have been developed that use reads mapped to one or more reference genomes to identify SNPs (Schloissnig et al. 2013; Lieberman et al. 2014) and gene copy-number variants (Greenblum et al. 2015; Zhu et al. 2015; Scholz et al. 2016) of microbial populations. These approaches have not been integrated together and/or made available as software. Recently, several software tools have been developed (Luo et al. 2015; Sahl et al. 2015) that use SNP patterns to phylogenetically type strains, but these methods do not capture the gene content of these organisms and may not be able to resolve strains in communities with high population heterogeneity. Additionally, existing methods do not provide comprehensive up-to-date genomic databases of bacterial species, thus limiting their utility across different environments. Assembly-based methods (Nielsen et al. 2014; Cleary et al. 2015) that seek to reconstruct microbial genomes without using reference genomes are a powerful alternative to assembly-free methods. However, these often require many samples, struggle to deconvolve closely related strains, or require manual inspection.

To address these issues, we developed the *Metagenomic Intra-species Diversity Analysis System (MIDAS)*, which is a computational pipeline that quantifies bacterial species abundance and intra-species genomic variation from shotgun metagenomes. Our method integrates many features (for a comparison to existing methods, see Supplemental Table S1) and leverages a comprehensive database of >30,000 reference genomes. Given a shotgun metagenome, *MIDAS* rapidly and automatically quantifies gene content and identifies SNPs in bacterial species, which is accurate for populations with a minimum of 1 and 10x sequencing coverage, respectively. These statistics enable quantitative analysis of bacterial populations within and between metagenomic samples.

To demonstrate the utility of this approach, we used *MIDAS* to conduct novel strain-level analyses on two datasets. First, we applied *MIDAS* to stool metagenomes from 98 Swedish mothers and their infants and used rare SNPs to track vertical transmission and temporal stability of strains in infants over the first year of life. Second, we used *MIDAS* to quantify gene content of prevalent bacterial species in 198 globally distributed marine metagenomes and identified significant intra-species population structure associated with geographic location and environmental variables. These analyses reveal striking structures in microbial communities that are missed when metagenomes are analyzed at a coarser taxonomic resolution.

## Results

### Identification of bacterial species with a consistent definition and efficient algorithm

To quantify strain-level genomic variation broadly and accurately, we built a comprehensive database of 31,007 high-quality bacterial reference genomes obtained from the Pathosystems Resource Integration Center (PATRIC) (Wattam et al. 2014). We accurately clustered these genomes into species groups to avoid inconsistent, erroneous, and incomplete annotations that afflict some microbial taxonomies (Mende et al. 2013), and to expand and improve upon previous efforts to systematically delineate bacterial species (Mende et al. 2013; Schloissnig et al. 2013; Varghese et al. 2015). Towards this goal, we hierarchically clustered reference genomes using the average pairwise percent identity across a panel of 30 universal genes (Figure 1a) that we selected from a panel of 112 candidates (Wu et al. 2013) (Supplemental Fig S1, Supplemental Table S2). We found that the best gene families for identifying bacterial species were less conserved and more widely distributed across the tree of life relative to other genes we tested (Supplemental Fig S2). For example, many ribosomal gene families were too conserved to differentiate closely related species (Supplemental Table S2). We applied a 96.5% nucleotide identity cutoff, which produced genome-clusters that were highly concordant with a gold standard definition of prokaryotic species based on 95% genome-wide average nucleotide identity (Konstantinidis et al. 2006; Richter and Rossello-Mora 2009) (Supplemental Table S3). Our procedure clustered the 31,007 bacterial genomes into 5,952 genome-clusters, representing distinct bacterial species (Supplemental Table S4-S5). We inferred the phylogenetic relationships of these species using a concatenated alignment of the 30 marker genes (Supplemental Fig S3). Because our algorithm uses a small set of highly informative marker genes, rather than genome-wide sequence comparisons, it will be efficient to update these genome-clusters as additional genomes are sequenced.

**Figure 1.**
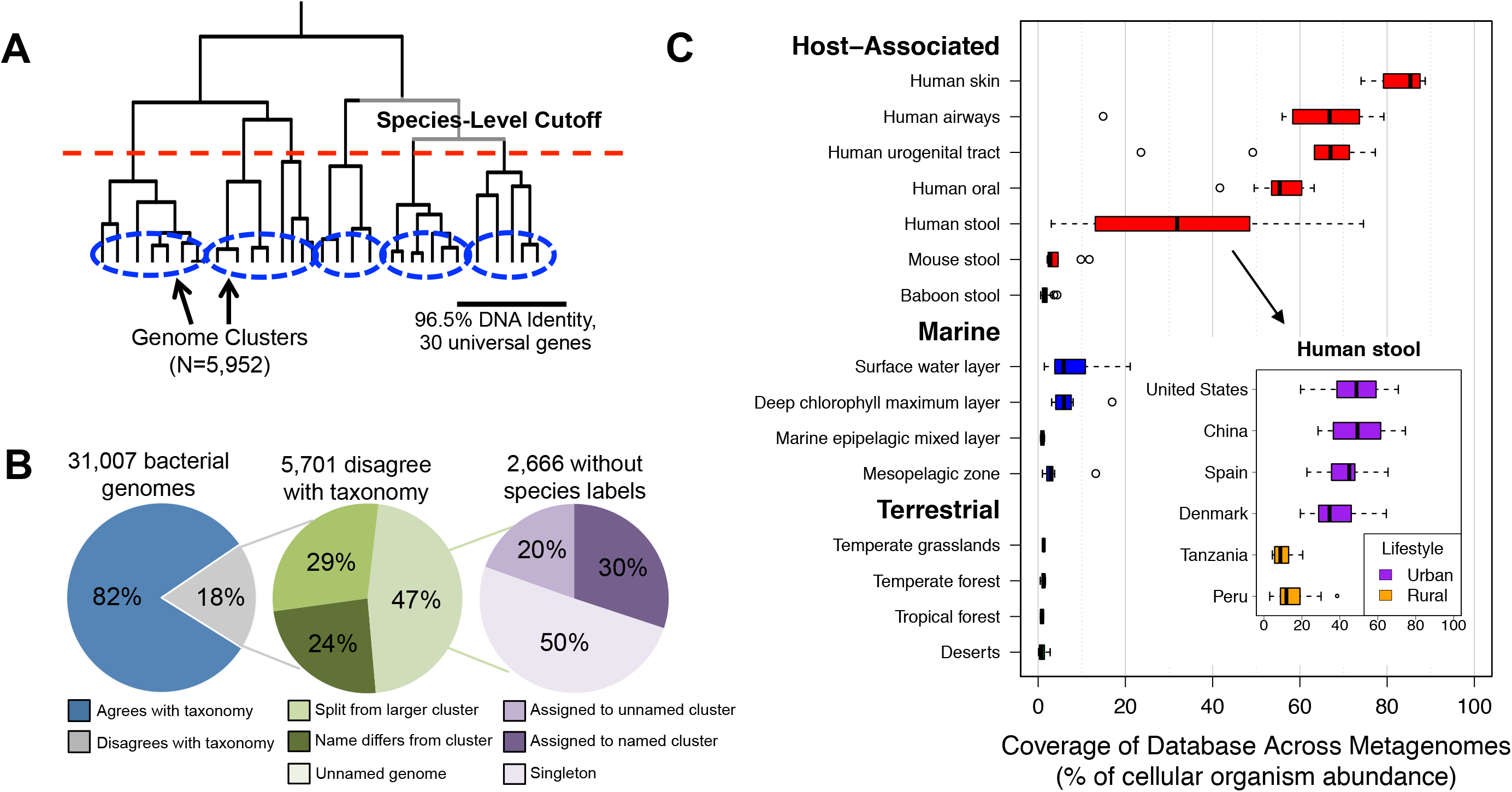
Construction of bacterial species database and its coverage of microbial communities across different environments. **A)** 31,007 genomes were hierarchically clustered based on the pairwise identity across a panel of 30 universal gene families. We identified 5,952 species groups by applying a 96.5% nucleotide identity cutoff across universal genes, which is equivalent to 95% identity genome wide. **B)** Concordance of genome-cluster names and annotated species names. Of the 31,007 genomes assigned to a genome-cluster, 5,701 (18%) disagreed with the consensus PATRIC taxonomic label of the genome-cluster. Most disagreements are due to genomes lacking annotation at the species level (47%). Other disagreements are because a genome was split from a larger cluster with the same name (29%) or assigned to a genome-cluster with a different name (24%). **C)** Coverage of the species database across metagenomes from host-associated, marine, and terrestrial environments. Coverage is defined as the percent (0 to 100%) of genomes from cellular organisms in a community that have a sequenced representative at the species level in the reference database. Inset panel shows the distribution of database coverage across human stool metagenomes from six countries and two host lifestyles.

The genome-clusters we identified often differed from the PATRIC taxonomic labels (Figure 1b). Our procedure clustered 2,666 genomes (8.6% of total) that had not been previously annotated at the species level and reassigned species labels for 3,035 genomes (9.8% of total) to either (i) group them with genomes that were not labeled as the same species in the reference taxonomy (N=1,380) or (ii) split them from genomes with the same label in the reference taxonomy (N=1,655). Supporting our species definitions, we found that the bacterial species we identified tended to have distinct functional repertoires, with only 0.05% of FIGfam protein families (Meyer et al. 2009) shared between genomes from different species on average compared to >80% for pairs of genomes from the same species. In previous work, Mende et al. conducted a similar procedure to cluster genomes into species groups and found that the majority of disagreements with the NCBI taxonomy were supported by the literature (Mende et al. 2013).

### Current reference genomes cover the majority of human-associated bacterial species and highlight novel diversity in other environments

We evaluated how comprehensively our reference database covers the abundance of species present in different environments, as this is a requirement for conducting reference-based strain-level analyses. Previous work has shown large gaps in diversity between sequenced reference genomes and environmental microorganisms (Wu et al. 2009). To explore how well current genome sequences cover diversity present in metagenomes from various environments, we developed a novel approach that estimates the proportion of microbial genomes (including archaea and eukaryotes, but excluding viruses) in a metagenome that contain a sequenced representative at the species level in a reference database (Methods). This proportion, which we call *database coverage*, indicates the degree to which species in a sample are known versus novel.

We applied this method to stool metagenomes from the Human Microbiome Project (HMP) and four other studies of the human gut (Supplemental Table S6). We found that our reference database of 5,952 bacterial species had high coverage of microbial communities from the human body (Figure 1c). This included high database coverage of samples from the skin (mean=83%), nasal cavity (mean=63%), urogenital tract (mean=62%), mouth (mean=55%), and gastrointestinal tract (mean=49%). The human gut communities with highest database coverage came from individuals in the United States, Europe, and China that live urban lifestyles, which is consistent with a previous report (Sunagawa et al. 2013). In contrast, gut microbiomes of individuals from Tanzania and Peru that live hunter-gatherer and agricultural lifestyles had much higher levels of novel species with no sequenced representative in our database. This finding extends the previous discoveries of elevated levels of novel genera (Schnorr et al. 2014) and functions (Rampelli et al. 2015) in the gut microbiome of African hunter-gatherers. Our analysis points to specific phylogenetic gaps in the set of currently sequenced bacterial genomes. Gut communities with lower database coverage (i.e., fewer species that have been sequenced) tended to have higher levels of several genera including *Coprococcus*, *Subdoligranulum*, *Dorea*, and *Blautia*, whereas well-characterized communities tended to have higher levels of the genus *Bacteroides* (Supplemental Fig S4). We conclude that there is a clear bias of genome sequencing to date towards species associated with hosts from industrialized countries.

In contrast to the human microbiome, a relatively small proportion of microbes present in other environments were captured by our reference database (Figure 1c and Supplemental Table S6). This included very low coverage for stool metagenomes from laboratory mice (mean=4.3%), which was surprising since mice are often used as a model system for studying the human microbiome. Coverage was also strikingly low in marine (means: surface water=8.2%, deep chlorophyll maximum layer=6.9%, subsurface epipelagic mixed layer=1.0%, mesopelagic zone=4.0%) and soil (means: desert=1.0%, forest=1.0%, grassland=1.3%, tundra=1.1%) environments. These estimates emphasize the massive gap that remains between the microbial diversity found in non-human environments and that represented by sequenced bacterial reference genomes. Strain-level analyses can still be performed for environments with low database coverage, but only for those species with sequenced representatives.

### An integrated pipeline for quantifying intra-species genomic variation from shotgun metagenomes

We developed *MIDAS*, a software tool that processes shotgun metagenomes to sensitively and automatically quantify species abundance and strain-level genomic variation for any of the bacterial species in our database (Figure 2a and Methods). *MIDAS* was designed to be fast, memory efficient, and to scale with the rapid increase in sequenced reference genomes (Supplemental Fig S5). Using a single CPU, *MIDAS* processes ∼5,000 reads per second and requires ∼3 gigabytes of RAM.

**Figure 2.**
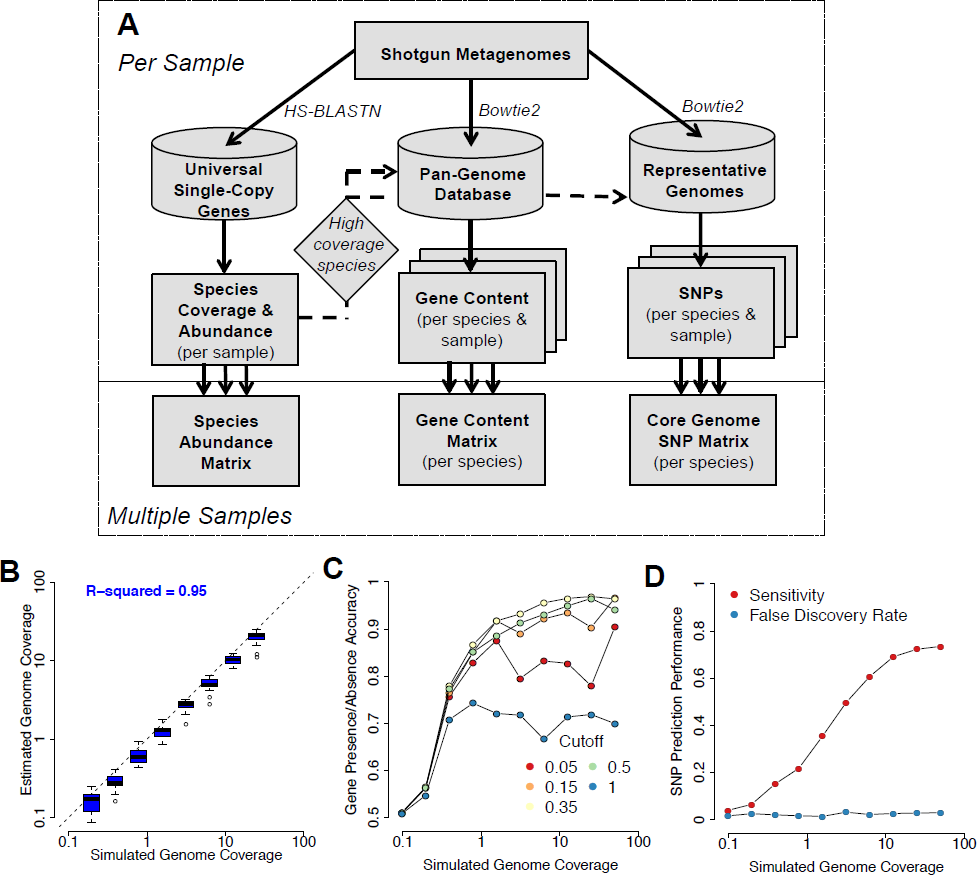
An integrated pipeline for profiling species abundance and strain-level genomic variation from metagenomes. **A)** The *MIDAS* analysis pipeline. Reads are first aligned to a database of universal-single-copy genes to estimate species coverage and relative abundance per sample. For species with sufficient coverage, reads are next aligned to a pan-genome database of genes to estimate gene coverage, copy-number, and presence/absence. Finally, reads are aligned to a representative genome database to detect SNPs in the core genome. The core genome is defined directly from the data by identifying high coverage regions across multiple metagenomic samples. **(B-D)** To evaluate performance for each component of MIDAS, we analyzed 20 mock metagenomes composed of 100-bp Illumina reads from microbial genome-sequencing projects. Each community contained 20 organisms with exponentially decreasing relative abundance. We tested the ability of MIDAS to estimate species coverage and to predict genes and SNPs present in the strains of the mock communities compared to the reference gene and genome databases. **B)** Species coverage is accurately estimated. Each boxplot indicates the distribution of estimated genome coverages across 20 mock communities for the top 8 most abundant species out of 20 analyzed. **C)** Gene presence/absence is accurately predicted when genome coverage is above 1x and a gene copy number cutoff of 0.35 is used. Accuracy = (Sensitivity + Specificity)/2; Sensitivity = (# genes correctly predicted as present)/(# total genes present); Specificity = (# genes correctly predicted as absent)/(# total genes absent). **D)** SNPs are detected with a low false discovery rate and good sensitivity when genome coverage is above 10x. Sensitivity = (# correctly called SNPs)/(# total SNPs); False Discovery Rate = (# incorrectly called SNPs)/(# called SNPs).

*MIDAS* first estimates the coverage and relative abundance of bacterial species by mapping reads to a database of universal single-copy gene families (Supplemental Table S7). Identifying species with sufficient coverage for gene content and SNP analyses directly from the shotgun metagenome enables automatic selection of individual species for variant quantification without any prior knowledge about a community’s composition, and it avoids computationally wasteful alignments to genes and genomes from sequenced organisms that are not present in a community.

To quantify the gene content of individual species in each metagenome, *MIDAS* maps reads to a pan-genome database. This database contains the set of non-redundant genes across all sequenced genomes from each species. It is generated on the fly to include only the subset of species with high sequencing coverage at universal single-copy genes in the metagenome being analyzed. The coverages of genes in the pan-genome database are normalized by the coverage of the universal single-copy gene families, yielding an estimated copy number of a gene per cell of a given species in each sample. Additionally, copy numbers are thresholded to predict gene presence-absence per sample.

To identify SNPs of individual species, *MIDAS* maps reads to a genome database. This database contains one representative genome sequence per species, and it only includes species with high sequencing coverage at universal single-copy genes in the metagenome being analyzed. Representative genomes are selected in order to maximize their sequence identity to all other genomes within the species. The core genome of each species is identified directly from the data using nucleotide positions in the representative genome that are at high coverage across multiple metagenomic samples (Supplemental Fig S6). SNPs are quantified along the entire core genome, including at sites that are variable between samples, but fixed within individual samples. Core genome SNPs are useful because they occur in all strains of a species and facilitate comparative analyses.

*MIDAS* was validated using 20 mock metagenomes that we created by pooling Illumina reads from completed genome sequencing projects (Supplemental Tables S8-9 and Methods). These libraries are expected to contain sequencing errors and other experimental artifacts found in real short-read sequencing data that might prevent accurate estimation of species abundance and strain-level genomic variation. Using this data, we found that *MIDAS* accurately estimated the relative abundance of bacterial species (r^2^=0.95), but slightly underestimated sequencing coverage (Figure 2b). *MIDAS* accurately predicted the presence or absence of genes in species present with at least 1 to 3x sequencing coverage (Figure 2c). Prediction accuracy was maximized at 0.96 for strains with >3x coverage when using a threshold equal to 0.35x the coverage of universal single-copy genes – lower thresholds resulted in lower specificity and higher thresholds resulted in lower sensitivity. *MIDAS* also called SNPs at a low false-discovery rate, but required between 5 to 10x coverage to identify the majority of SNPs present (Figure 2d).

### Species and strain-resolved analyses shed light on vertical transmission of human gut microbiota

We hypothesized that the large numbers of SNPs that *MIDAS* can identify from individual metagenomes could be leveraged to detect bacterial strains unique to a host and transmission of strains between mothers and their infants. An understanding of vertical transmission is critical for determining the extent to which the microbiome – and by extension microbiome-mediated phenotypes – are inherited. A recent study found significant overlap in species between Swedish mothers and their infants over the first year of life (Backhed et al. 2015). A large cohort study of UK twins estimated that abundances of many microbial taxa are heritable (Goodrich et al. 2016). Neither study examined whether strains are vertically transmitted, and recent work has shown that species-level analyses alone are insufficient to resolve transmission events (Li et al. 2016). Transmission of specific taxonomic groups has been resolved using multilocus sequence typing of infant stool plus mother’s breast milk (Martin et al. 2012) or stool (Makino et al. 2013). Other studies have examined the development of the infant gut microbiome, including at the strain level (Luo et al. 2015), but did not assess vertical transmission. Thus, the extent and timescale of vertical transmission and the stability of transmitted stains are currently not well established.

To quantify strain transmission from mother to infant, we applied *MIDAS* to the Backhed et al. stool metagenomes from 98 mothers and their infants at 4 days, 4 months, and 12 months after birth (Backhed et al. 2015). We found that bacterial species alpha diversity was lowest in newborns and increased over time, species beta diversity was highest in newborns and decreased over time, and samples clustered by host age based on Bray-Curtis dissimilarity between species relative abundance profiles (Figure 3a and Supplemental Fig S7). Compared to infants, mothers had more diverse microbiomes that tended to harbor more unshared (i.e., unique to host) species (77% versus 48%, T-test P<2.2e-16). Despite this, we found a large number of shared species between infants and their mothers, which increased over time as diversity increased in the infants (Figure 3b) and did not strongly depend on birth mode (P=0.52) or breast-feeding at any stage (P-values = 0.04, 0.04, 0.33 at 4 days, 4 months, and 12 months). These species-level trends agree with the results of the original study that used different methods to identify species (Backhed et al. 2015). Surprisingly, we found nearly as many shared species between permuted mother-infant pairs where vertical transmission did not occur (Figure 3c), suggesting that increased similarity of species in a mother and her infant over its first year is unlikely the result of direct transmission.

**Figure 3.**
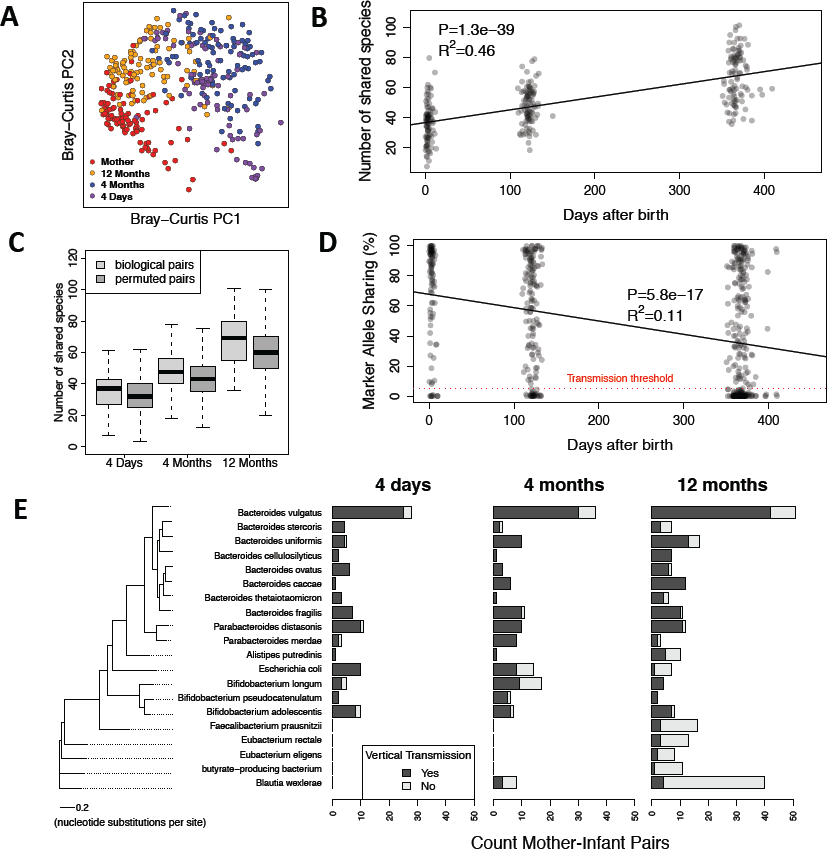
An increase in shared species but decrease in shared strains over time between stool metagenomes from mothers and their infants. **A)** Principal coordinate analysis of Bray-Curtis dissimilarity between species relative abundance profiles of stool samples from mothers and infants at 4 days, 4 months, and 12 months following birth. Species composition of infant microbiomes is most similar to mothers at 12 months. **B)** The number of shared species increases over time between mothers and their own infants. **C)** This pattern for biological mother-infant pairs is similar to that of unrelated mothers and infants (permuted pairs). **D)** In contrast, marker allele sharing decreases over time between mothers and their infants for shared species with >10x sequencing coverage, indicating highest strain similarity at 4 days. Allele sharing is defined as the percent of marker alleles in the mother that are found in the infant. The horizontal red line indicates the 5% marker allele threshold used for defining vertical transmission events. **E)** Vertical transmissions for 20 bacterial species across mother-infant pairs at three time points. A vertical transmission is defined as >5% marker allele sharing between mother and infant. The phylogenetic tree on the left is constructed based on a concatenated DNA alignment of 30 universal genes (Supplemental Fig S3). The tree shows that species with more vertical transmission are phylogenetically clustered.

To detect transmission of gut microbiota from mother to infant with high specificity and sensitivity, we developed a novel approach that uses SNPs output by *MIDAS* (Methods). First we identified species shared between mothers and their infants with >10x sequencing coverage, which is required for sensitive detection of SNPs (Figure 2d). Next, we identified rare SNPs within these species that were private to strains found in a mother and her infant. We refer to these SNPs as *marker alleles* because they serve as a marker for individual strains. To detect whether a transmission has occurred for a species, we quantified the percent of marker alleles found in a mother that were shared with her infant.

To validate that marker alleles could be used to track strains between hosts, we applied our method to stool metagenomes of healthy adults from the Human Microbiome Project (HMP) (Methods). As a positive control, we compared marker alleles of species between metagenomes from the same individual at the same time point (technical replicates), and as a negative control, we compared marker alleles of species between metagenomes from different unrelated individuals (non-replicates). As expected, we found high allele sharing (mean=79.5%) between technical replicates and low allele sharing between non-replicates (mean=1.01%) (Supplemental Fig S8). The fact that allele sharing was <100% in the technical replicates and >0% in the non-replicates is likely due to a combination of factors, including read sampling variation, small sample sizes, and sequencing errors. For example, marker alleles may be found in other individuals when sample sizes are increased. To define a transmission event, we selected a marker allele sharing cutoff of 5%, which clearly separated the positive and negative controls (sensitivity=99.8%, specificity=96.6%). High sensitivity and specificity was consistently observed across species we tested (Supplemental Table S10).

Strikingly, we found that marker alleles were commonly shared between mothers and vaginally born infants 4 days after birth (Figure 3d-e). There were no species present with >10x coverage in 15 C-section born infants and their mothers to assess transmission in these individuals. This likely reflects lower vertical transmission of the mother’s gut microbes, but we cannot directly test that hypothesis with the available data. On average 72% of marker alleles present in mother strains were found in vaginally born newborns, which was only slightly less than the level of allele sharing observed from our positive control. Furthermore, out of the 111 high-coverage species present in mothers and newborns, 101 (91%) had greater than 5% marker allele sharing, indicating extensive vertical transmission of gut microbiota shortly after birth. Commonly transmitted species included *Bacteroides vulgatus* (25/28 mother-infant pairs with >5% marker allele sharing), *Parabacteroides distasonis* (10/11), *Bifidobacterium adolescentis* (8/10), and *Escherichia coli* (10/10) (Figure 3d). These are fairly different from the taxa with heritable abundances in the UK twin study (Goodrich et al. 2016) (see Discussion).

While we detected high strain similarity 4 days after birth, mother and infant strains significantly differed over time. Comparing strain-level SNPs in 4-month and 12-month infants to their mothers, we observed a sharp decrease in marker allele sharing and transmission rates (Figure 3d). Across all species, transmission rates decreased from 91% at 4 days (101/111 shared species with >5% marker allele sharing), to 80% at 4 months (131/163), to 55% at 12 months (172/313). C-section born infants tended to have fewer vertically strains transmitted strains compared to vaginally born infants at four months (Chi-square P=5e-8, 3/14 versus 128/149 shared species with >5% marker allele sharing) and to a lesser extent at 12 months (Chi-square P=0.06, 13/34 versus 159/279). This trend was in stark contrast to what we observed at the species level, where there was an increase in the number of shared species over time and an increase in species-level compositional similarity. Thus, while the species level composition of mothers and infants converged over time, the strain level composition actually diverged.

We hypothesized that transmission rates decreased over time due to late colonization of the infant gut by new strains from the environment and/or other hosts. If this were the case, then we would expect that (i) infant strains that were distinct from the mother at 12 months had low abundance in the infant at earlier stages and (ii) strains transmitted from the mother at 4 days persisted in the infant gut over one year. Supporting our hypothesis, we found that the abundance of a species at 4 days was predictive of whether the strain of that species was transmitted from the mother (Figure 4a). Specifically, strains with low abundance at 4 days but high abundance at 12 months tended to be distinct from strains found in the mother. In contrast, strains with high abundance at 4 days and high abundance at 12 months were similar to strains found in the mother. Also supporting our hypothesis, we found that the vast majority of strains that were transmitted from the mother at 4 days persisted in the infants at 4 months (49/54 mother-infant pairs with >5% marker allele sharing) and at 12 months (47/51). Because the mother’s stool was only sequenced at 4 days after birth, we cannot rule out the possibility that late colonizing strains came from the mother’s gut but were not detected at the time of initial sampling. To address this issue, we quantified the temporal stability of strains in 157 healthy adults from the HMP over a time period of 300-400 days (Supplemental Fig S9). We found high marker allele sharing (mean=77.0%) and transmission rates (96.2%), which suggests that maternal strains may be quite stable over time, and agrees with previous work (Faith et al. 2013; Schloissnig et al. 2013). Together these results suggest that bacteria from sources other than the mother’s gut increasingly colonize the infant gut over time.

**Figure 4.**
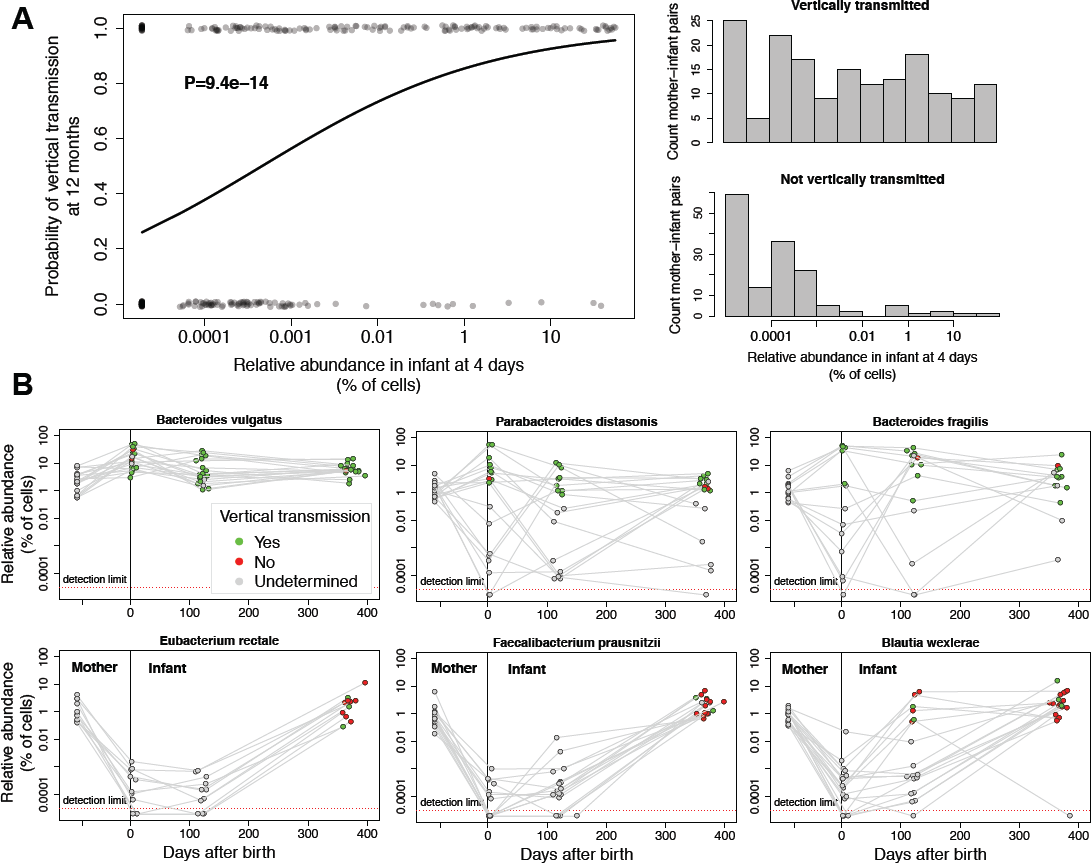
Strains that colonize the infant gut late in the first year are rarely found in the mother. **A)** Vertical transmission of strains at 12 months versus their relative abundance at 4 days. Each data point indicates a shared species found in a mother and her infant. The vertical axis indicates whether a strain of the species was transmitted from the mother (y=1) or not (y=0). The curve is a logistic regression fitted to data points. Histograms indicate the distribution of relative abundance at 4 days for strains that were transmitted and not transmitted. Strains with high relative abundance in infants at 4 days are frequently transmitted from the mother. Strains that appear in the infant at later time points (i.e. low abundance at 4 days) are rarely transmitted from the mother. **B)** Examples of early colonizing species that are frequently transmitted vertically ( *Bacteroides vulgatus, Parabacteroides distasonis*, and *Bacteroides fragilis*) and late colonizing species that are rarely transmitted vertically ( *Eubacterium rectale, Faecalibacterium prausnitzii*, and *Blautia wexlerae*). Points are colored based on vertical transmission. Gray indicates there was insufficient sequencing coverage to quantify SNPs and determine whether the strain was transmitted from the mother or not.

We found that vertical transmission rates varied for different taxonomic groups of bacteria. At one year after birth, the class Bacteroidia was enriched in vertical transmission events (Chi-square P=2.56e-18), whereas the class Clostridia was depleted (Chi-square P=1.43e-22). Similar results were observed at other taxonomic levels (Supplemental Table S11). Examples of early colonizing Bacteroidia included *Bacteroides vulgatus*, *Bacteroides uniformis*, and *Bacteroides ovatus* and examples of late colonizing Clostridia included *Blautia wexlerae, Faecalibacterium prausnitzii*, *Eubacterium rectale*, and *Ruminococcus bromii* (Figure 4b). The fact that Clostridia were rarely transmitted from mother to infant may be due to the ability of members of this group to form spores and survive outside of the host for longer periods of time (Browne et al. 2016). Together, these results highlight differences in the inheritance of gut microbiota that may be linked to distinct modes and timing of transmission between hosts.

### Global strain-level geography of prevalent marine bacteria

Many bacterial species are distributed widely across the world’s oceans (Sunagawa et al. 2015). Yet genomes of a given species sampled nearby each other can differ significantly in their gene content (Kashtan et al. 2014). To explore the extent of population structure across different marine bacterial species on a global scale, we used *MIDAS* to quantify gene content for prevalent species in 198 marine metagenomes from 66 stations along the *Tara* Oceans expedition (Sunagawa et al. 2015). Since we found that our database had relatively low coverage of the cellular organisms present ocean samples (Figure 1c), we first estimated relative abundance and coverage of bacterial species in each metagenome to identify 30 species where gene content could be reliably estimated (coverage >3x across a high percentage of samples). Among these species were several members of the genera *Pelagibacter*, *Alteromonas*, *Synechococcus*, and *Marinobacter*, a large group of closely related *Prochlorococcus* species, and several unnamed *Alphproteobacteria* species (Figure 5a). Reference pangenome sizes for these species ranged from 1,047 and 1,311 genes in the streamlined genomes of SAR406 and SAR86 (each with 1 genome) to 6,427 genes in the largest *Prochlorococcus* genome cluster (N=26 genomes) and 7,819 genes for *Alteromonas macleodii* (N=4 genomes).

**Figure 5.**
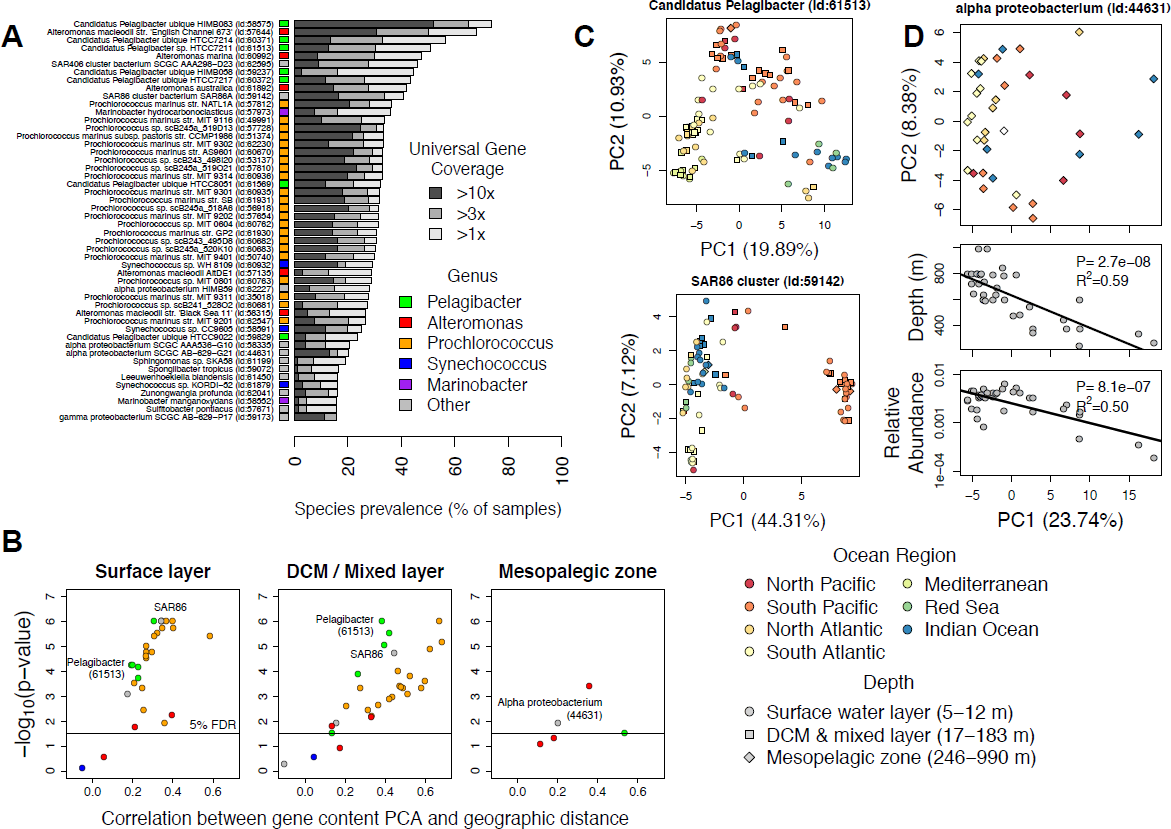
Gene content and geography are correlated for many marine bacteria. **A)** Prevalence of 50 bacterial species across 198 ocean metagenomes. Species identifiers are indicated in parenthesis. Many marine species have sufficient sequencing depth and prevalence for population genetic analyses. **B)** Correlation of PCA and geographic distance between pairs of samples for different marine species. PCA distance was calculated using the Euclidian distance between PC1 and PC2 of the gene presence/absence matrix. Geographic distance was calculated using the great-circle distance between sampling locations. Correlations and p-values were computed using the Mantel test with 1 million permutations. Only one metagenome per location was included in the tests. The population structure of marine bacteria, based on the first two principal components of gene content, is correlated with geography for many species of bacteria. **C)***Candidatus Pelagibacter* and SAR86 are examples of marine bacterial species geographically structured populations based on gene presence/absence. Each point indicates a bacterial population from a different sample. Point colors indicate the marine region. Point shapes indicate the ocean depth. Samples tended to cluster together based on ocean region, not ocean depth. **D)** *Alpha proteobacterium* is an example of a species present in the mesopelagic zone where depth is the major driver of population structure.

We discovered extensive variability of gene content for these prevalent species of marine bacteria across the ocean metagenomes (Supplemental Table S12). Across all species, we found an average of 318 genes that differed between samples, ranging from 144 genes in *SAR86* to 700 in *Alteromonas marina*. We next quantified the percent of genes that were different between samples using the Jaccard index and found that on average 19% of genes differed between samples. This level of genomic variability was higher than the 13% recently reported for human gut communities (Zhu et al. 2015), although this may be due to methodological differences. Regardless, our estimate of 19% is almost certainly an underestimate of the true level of gene content variation between populations, because *MIDAS* cannot measure the variation of genes that are present in strains but absent from sequenced reference genomes.

To explore how this variation correlated with geography and sampling depth, we conducted a principal component analysis (PCA) of gene content for each bacterial species, as has been done to study the geographic structure of human populations (Novembre et al. 2008). Strikingly, we found that the populations of many species clustered together by ocean region based on the first two principal components of gene content, regardless of sampling depth (Figure 5b). For example, populations of *Pelagibacter* (species id: 61513) formed three discrete clusters corresponding to the Mediterranean Sea, South Atlantic Ocean, and South Pacific Ocean, and each cluster contained samples from different water layers. Similar results were obtained for many other species, including *Prochlorococcus* (species id: 57810) and *SAR86* (species id: 59142) (Figure 5c). We found that the population structure of the 30 marine bacteria was highly consistent, regardless of the percent identity threshold used for defining pan-genome gene families (75-99% identity) (Supplemental Fig S10-11).

To evaluate the extent of gene content biogeography across species, we computed the correlation between PCA distances and geographic distances (Methods) and found significant distance-decay in gene content for the majority of species tested, including all 18 *Prochlorococcus* species (Figure 5b). Furthermore, this pattern was observed both in samples from the surface water layer and the deep chlorophyll maximum layer. A previous study found season to be a major driver of biodiversity patterns in the global ocean (Ladau et al. 2013). To explore whether season or other environmental variables were associated with strain-level population structure, we compared correlations of the first principal component of gene content (PC1) with geography and environmental variables (Supplemental Fig S12). For 20/30 species tested, longitude (17/30) or latitude (3/30) was the strongest predictor of gene content, and each explained a significant proportion of gene content variation (22% and 8% on average). In contrast, day length (an indicator of season) explained relatively little variation (4% on average) and was the most predictive covariate for only one *Prochlorococcus* species (species id: 60683).

A few species showed relatively little geographic structure. Instead they had gene content variation that correlated with depth or marine layer. The most striking example of this was an unnamed *Alphproteobacteria* species (species id: 44631) which contained two genomes in our database obtained via single-cell sequencing (Stepanauskas 2012). This species was predominantly found in the mesopelagic layer (below 200m) and increased in relative abundance with decreasing depth (Figure 5c). Looking only at mesopelagic samples, we found that the first principal component of gene content (PC1) was strongly correlated with depth (R^2^=0.59) suggesting little mixing of strains of the *Alphproteobacteria* species across depths. We identified 266 genes positively correlated with depth and 316 genes that were negatively correlated with depth (FDR-corrected Spearman p-value < 0.01, Supplemental table S13). This could indicate that the populations at different depths contain genes for adaptation to the range of temperatures and nutrients across which this species is found. When we included samples from all marine layers, we found that samples from the mesopelagic and epipelagic zone clustered based on gene content and there was still a strong correlation between PC1 and depth (R^2^=0.57) (Supplemental Fig S13).

Together, our results expand upon and even contradict patterns of marine bacterial biogeography observed at the species level. In particular, gene content analysis reveals that abundant and prevalent species are not ubiquitous at the strain level. Instead they show significant structure across geographic regions, even though sampling location is not a strong predictor of species relative abundance (Sunagawa et al. 2015).

## Discussion

We developed *MIDAS*, an integrated computational pipeline that quantifies bacterial strain-level gene content and SNPs, as well as species abundance, from shotgun metagenomes. By coupling fast taxonomic profiling via a panel of universal-single-copy genes with sensitive pan-genome and whole-genome alignment, *MIDAS* can efficiently and automatically compare hundreds of metagenomes to >30,000 reference genomes to identify genetic variants present in the strains of each sample. Our publicly available software and data resources will enable researchers to conduct large-scale population genetic analysis of metagenomes.

This first version of *MIDAS* has several limitations. Since it currently relies on bacterial reference genomes, *MIDAS* cannot quantify variation for novel species, novel genes, or known species from other groups of microbes (e.g. archaea, eukaryotes, and viruses). To accurately quantify strain-level gene content and SNPs, *MIDAS* requires greater than 1x and 10x sequencing coverage, respectively. This biases analyses towards the most abundant and prevalent species in an environment. *MIDAS* was nonetheless able to capture the majority of microbial species abundance across human body sites, making it well suited for uncovering strain-level variation of human-associated bacteria. In contrast, other environments appeared to be dominated by microbes missing from our reference database of bacterial genomes sequenced to date. For this reason, it will be important to update the database in the future as the number (Land et al. 2015) and diversity (Wu et al. 2009) of microbial reference genomes continues to rapidly grow, as new experimental (Rinke et al. 2013) and computational (Nielsen et al. 2014) approaches uncover genome sequences of uncultured microbes. It will also be useful to incorporate genomes from other domains of life. Based on the design of our database and algorithm, *MIDAS* should scale with this growth of reference data.

To illustrate the utility of *MIDAS*, we analyzed stool metagenomes from 98 mothers and their infants over one year and used rare SNPs (i.e. marker alleles) to track transmission of strains between hosts. Based on this analysis, we found extensive vertical transmission of specific early colonizing bacteria shortly after birth, which largely persisted in the infant for one year. In contrast, we found that late colonizing bacteria were often distinct from the mother at the strain level, likely originating from the environment and/or other hosts. Additionally we found that certain taxonomic groups, like Bacteroidia, tend to be vertically transmitted, while others, like Clostridia do not, which suggests that only part of the gut microbiome may be inherited. When the same metagenomes were analyzed at the species level, these patterns of transmission were missed, and a false signal of increasing transmission over time was detected due to convergence of the infant microbiome towards a more diverse and adult-like species profile after weaning.

The bacterial taxa that tended to be transmitted from mother to infant in our analysis differ from the taxa whose abundances were found to be heritable in a previous study of UK twins (Goodrich et al. 2016). For example, we estimated low transmission rates for strains of *Blautia* (Supplemental Table S11), but Goodrich et al. found that the abundance of *Blautia* is highly heritable. Conversely, we estimated high transmission rates for strains of *Bacteroides*, but this was one of the genera whose abundance was least heritable in the UK twins. These differences between studies reveal some important distinctions. First, heritability of taxon abundance does not necessarily imply vertical transmission. Rather, twins and other related individuals may share a propensity to retain similar levels of taxa that they both acquire from a variety of sources (i.e., different strains). On the other hand, strains vertically transmitted at birth should result in heritability of presence if they are retained. But they could be lost as the infant ages, or their abundances may not be heritable even if their presence is. Finally, human genetics may not explain inter-individual differences in abundance for taxa, such as *Bacteroides*, that are highly abundant and prevalent, even if they are vertically transmitted.

Our analysis of mother-infant strain sharing leaves a few questions unanswered. One intriguing issue is the source of the strains that colonize the infant but are not present in the mother’s stool microbiome at 4 days after birth. It is possible that some strains colonize the mother’s gut later in the year and are then passed along to the infant, though this is unlikely based on the temporal stability of strains in the adult microbiome. The new strains could also derive from other sites on the mother’s body, such as skin and breast milk, other people, or the environment. One caveat of our analysis is that we did not distinguish which strains were transmitted to the infant from the mother in cases where mothers harbored multiple strains. Instead, we treated the transmission events as binary, whereby a transmission was defined as at least one strain being transmitted. It would be interesting to explore transmission as a quantitative variable in future work, including elucidating how the strain composition and genetic diversity of bacterial populations change as they are passed from mother to offspring and potentially undergo bottlenecks and selection.

To explore bacterial population structure using gene content, we applied *MIDAS* to ocean samples from the *Tara* expedition. We found a number of prevalent and abundant bacterial species, which shows that our method can be applied to different environments, despite low database coverage. Based on these results, we found that the gene content of many species in the epipelagic water layer (0-200m) was structured geographically. This contrasts with previous work at the species level, which found that depth and temperature were the strongest predictors of community structure (Sunagawa et al. 2015). However, the gene content of other species found in the mesopelagic layer (200-1000m) were structured by depth. Future work is needed to understand the extent to which these gene-level patterns are driven by adaptation to different environments in the ocean, or due to neutral processes, like genetic drift and/or migration.

Microbiome research is in an era where metagenome-wide analyses can now pinpoint individual strains and genes that differ in presence or abundance between samples. Importantly, this level of resolution is not only revealing associations that are missed by analyses conducted at higher taxonomic levels, but also patterns that oppose those inferred from species abundance distributions. A striking example is our discovery that infants share more gut bacterial strains with their mothers at birth than later in their first year of life, despite the fact that the species composition of their microbiomes are becoming more similar as the infant ages. Without conducting a strain-level genomic analysis, one might incorrectly infer that vertical transmission of the gut microbiome is constant or increasing during the first year of life. Similarly, the high level of gene copy number variation that we observe in *Tara* oceans bacteria and its strong correlation with marine region in surface waters emphasizes functionally important differences in strains across global oceans that are missed when metagenomes are analyzed at the species level. Furthermore, it is clear that gene content of even the most prevalent and abundant marine bacterial species cannot be accurately inferred from the currently limited number of sequenced genomes for this environment. The same is true for laboratory mice, humans from non-industrialized countries, and soil. These results point to specific phylogenetic groups and environments that are highest priority for additional genome sequencing, culturing attempts, and functional assays.

## Materials and Methods

### Sequence based identification of bacterial species

We developed a procedure to cluster bacterial genomes into species groups based on the pairwise percent identity across a set of universal gene families, which was inspired by previous work (Mende et al. 2013). We began with 33,252 prokaryotic genomes downloaded from PATRIC (Wattam et al. 2014) in March 2015. Next, we used HMMER3 (Eddy 2011) with an E-value threshold ≤1e-5 to identify protein homologs of 112 bacterial universal gene families (Wu et al. 2013) across the genomes. The HMMER3 search took too long for two gene families (B000042, B000044), which were dropped. When there were multiple homologs of a gene family identified in a genome, we took the homolog with the lowest E-value. We filtered out low quality genomes with fewer than 100 universal genes identified (N=1,837) or with greater than 1,000 contigs (N=618), which left 31,007 high-quality genomes (Supplemental Table S5).

Next, we used BLASTN (Altschul et al. 1990) to perform sequence alignment of each gene family between all high-quality genomes. We filtered out local alignments where either the query or target was covered by <70% of its length. We converted percent identities to distances using the formula: *D_ab_* = (100 − *P_ab_*)/100, where *P_ab_* was the percent identity of a gene between genomes *a* and *b*. This resulted in an undirected graph for each marker gene family where nodes were genomes and edges were distances. We performed average-linkage hierarchical clustering for each graph using the program MC-UPGMA (Loewenstein et al. 2008). The output of MC-UPGMA is a tree, which we cut at different distance thresholds (0.01 to 0.10). Each cut of the tree yielded a set of genome-clusters.

For validation, we compared each set of genome-clusters to average nucleotide identity (ANI), which is considered to be a gold standard for delineating prokaryotic species (Konstantinidis et al. 2006; Richter and Rossello-Mora 2009) but was too computationally intensive to compute for all genome-pairs. Specifically, we used the procedure described by Richter and Rossello-Mora (Richter and Rossello-Mora 2009) to compute ANI for >18,000 genome-pairs and labeled pairs of genomes with ANI ≥ 95% as members of the same species and pairs of genomes with ANI < 95% as members of different species. We compared these labels to our genome-clusters and classified each genome-pair into one of the following categories: *True positive*: a clustered genome-pair with ANI ≥ 95%; *False positive*: a clustered genome-pair with ANI < 95%; *False negative*: a split genome-pair with ANI ≥ 95%; *True negative*: a split genome-pair with ANI < 95%. Using these classifications we calculated the true positive rate (TPR), precision (PPV), and F1-score for each set of genome-clusters corresponding to 90-99% identity between pairs of genomes for a given marker gene (Supplemental Table S2).

Based on this evaluation, we identified a subset of 30 gene families that produced genome-clusters that were in agreement with ANI, all with maximum F1-score > 0.93 across thresholds. To increase clustering performance, we took the average pairwise distances across these 30 gene-families and used these new distances to re-cluster genomes using MC-UPGMA (Supplemental Table S3). We found that a distance cutoff of 0.035 (96.5% nucleotide identity) maximized the F1-score at 0.98 and resulted in 5,952 genome-clusters (Supplemental Table S3). Each genome-cluster was annotated by the most common PATRIC Latin name within the cluster (Supplemental Table S4).

### Genomic database construction

Genome-clusters (i.e. bacterial species) were leveraged in order to compile a comprehensive genomic data resource used by *MIDAS*. First, we identified a representative genome from each species to use for detecting core-genome SNPs. Each representative genome was chosen in order to maximize its average nucleotide identity at the 30 universal genes (Supplemental Table S2) to other members of the species. Next, we build a database of 15 universal single-copy gene families (Supplemental Table S7) to use for estimating the abundance of the species from a shotgun metagenome. Gene families were selected based on their ability to accurately recruit metagenomic reads as well as being universal and single-copy. Many of the 30 gene families for clustering genomes into species and the 15 gene families for quantifying species abundance from metagenomes were different and were selected using distinct criteria. Next, we used USEARCH (Edgar 2010) to identify the set of unique genes at 99% identity across all genomes within each species, which are used by *MIDAS* for metagenomic pan-genome profiling. This procedure clustered 116,978,184 genes from the 31,007 genomes into 31,840,245 gene families. We further clustered these genes at different levels of sequence identity (75-95% DNA identity) in order to identity *de novo* gene families of varying size and diversity for downstream analyses. Functional annotations for all genes were obtained from PATRIC and include FIGfams (Meyer et al. 2009), Gene Ontology (Consortium 2000), and KEGG Pathways (Kanehisa and Goto 2000).

### Species abundance estimation

*MIDAS* uses reads mapped to 15 universal single-copy gene families to estimate the abundance of the 5,952 bacterial species from a shotgun metagenome. These 15 gene families were selected from a set of 112 phylogenetically informative bacterial gene families (Wu et al. 2013) for their ability to accurately recruit metagenomic reads to the correct species. To evaluate how informative different gene families are for estimation of abundance, we simulated one hundred 100-bp reads from each of the 112 gene families in each of the 5,952 species and used HS-BLASTN (Chen et al. 2015) to map these reads back to a database that contained the full length gene sequences. To simulate the presence of novel species and strains, we discarded alignments between reads and reference sequences from the same species. Each read was assigned to a species based on its top hit. Recruitment performance was measured using the F1-score. Based on this experiment, we identified 15 universal single-copy gene families that were best able to accurately assign the species from which metagenomic reads derived. Additionally, we identified the optimal percent identity cutoffs for mapping reads to the database, which ranged from 94.5% to 98.00% identity depending on the gene family (Supplemental Table S7).

To perform taxonomic profiling, *MIDAS* aligns reads to the database of 15 universal gene families with HS-BLASTN, discards local alignments that cover <70% of the read or alignments that fail to satisfy the gene-specific species-level percent identity cutoffs, and assigns each uniquely-mapped read to a species according to its best-hit. *MIDAS* assigns non-uniquely mapped reads (i.e. identical alignment scores to genes from >1 species) using probabilities estimated from uniquely mapped reads. These mapped reads are used to estimate the coverage and relative abundance of each species.

### Gene content estimation

To estimate gene content, *MIDAS* first uses the species abundance profile to identify bacterial species with sufficient coverage (e.g. >1x). A pan-genome database is dynamically built, which contains a set of non-redundant genes from these species. We used a 99% sequence identity threshold to cluster similar genes such that any two genes that are <99% similar were classified as distinct genes. Bowtie2 (Langmead and Salzberg 2012) is used to locally map reads from the metagenome against the pan-genome database. Each read is mapped a single time according to its best hit, and reads with an insufficient mapping percent identity (default=94%), alignment coverage (default=70%), mapping quality (default=20), or sequence quality (default=20) are discarded.

Mapped reads are used to compute the coverage of the genes clustered at 99% identity. Since the 99% identity may result in many very similar gene families, *MIDAS* gives the option of further clustering the gene families at lower sequence identities ranging from 75% to 95%. Aggregating enables quantification of gene families of varying size and diversity, while maintaining mapping speed and sensitivity.

To estimate gene copy numbers in a bacterial population, gene coverages are normalized by the median coverage across the 15 universal single-copy gene families (Supplemental Table S7). Copy-number values are thresholded to produce gene presence/absence calls. *MIDAS* merges these results across multiple metagenomic samples to produce gene content matrices for all species, which facilitate comparative analyses across genes and metagenomic samples.

### Identifying core genome SNPs

To estimate core genome SNPs, *MIDAS* first uses the species abundance profile to identify species with sufficient coverage (e.g. >10x). A representative genome database is dynamically built, which contains a single genome per species that meets the coverage requirement. The representative genome is a single genome chosen that has the greatest nucleotide identity, on average, to other members of the species. Only a single genome is needed for identifying the core genome, because this region should be present in all strains of a species. Bowtie2 is used to globally map reads to the representative genome database. Each read is mapped a single time according to its best hit, and reads with an insufficient mapping percent identity (default=94%), alignment coverage (default=70%), mapping quality (default=20), or sequence quality (default=20) are discarded. Additionally, bases with low sequence quality scores are discarded (default=30). Samtools (Li et al. 2009) is used to generate a pileup of nucleotides at each genomic position and generate VCF files. VCF files are parsed to generate output files that report nucleotide variation statistics at all genomic sites. To identify the core genome of a species, *MIDAS* uses the output from multiple metagenomic samples to identify regions at consistently high coverage (e.g. >10x coverage in 95% of samples) (Supplemental Fig S6). *MIDAS* then produces core genome SNP matrices for all species, which facilitate comparative analyses of nucleotide variation across genomic sites and metagenomic samples. *MIDAS* also gives the option of outputting all SNPs, including those that are not in the core genome.

### Shotgun simulations and validation of *MIDAS* output

To validate *MIDAS* we designed a series of realistic metagenomic simulations using reads from completed genome-sequencing projects deposited in the NCBI Sequence Read Archive (Leinonen et al. 2011) which we identified using the SRAdb tool (Yuelin Zhu 2013). We used this data to construct 20 mock metagenomes, which each contained 100-bp Illumina reads from 20 randomly selected bacterial genome projects (Supplemental Tables S8-9). We only selected genome projects that corresponded to one of the 31,007 genomes present in our reference database, and we used only one genome project per selected species. We simulated libraries that contained 100x total genome coverage. The relative abundances of the 20 genomes were exponentially distributed in each simulation (50%, 25%, 12%, 6.5% etc.).

We compared the output of *MIDAS* to the known species abundance, gene content, and SNPs in the simulated communities. To evaluate the accuracy of species abundance estimation we compared the expected relative abundance and coverage to the simulated relative abundance and coverage. To evaluate the accuracy of gene content estimation, we ran *MIDAS* to estimate the copy-number of genes in the pan-genome of each species in each simulation. We applied a cutoff to these values to predict gene presence/absence. True positives (TP) were present genes predicted as present, false positives (FP) were absent genes predicted as present, true negatives (TN) were absent genes predicted as absent, and false negatives (FN) were present genes predicted as absent. Performance was measured across a range of copy-number cutoffs using balanced accuracy: (TPR+TNR)/2, where TPR=TP/(TP+ FN) and TNR=TN/(TN+FP). To evaluate the accuracy of core genome SNPs, we ran *MIDAS* to estimate the frequency of nucleotide variants in the representative genome of each species in each simulation. We predicted SNPs using the consensus allele at each genomic position. True SNPs were identified by comparing genomes in the simulations to the representative genomes used for read mapping with the program MUMmer (Stefan Kurtz 2004), which identified 3,971,528 total true SNPs. True positives were correctly called SNPs, false positives were incorrectly called SNPs, and false negatives were SNPs that were not called due to insufficient coverage. We compared predicted SNPs to true SNPs and measured performance using the true positive rate (TP/TP+FN) and precision (TP/TP+FP).

### Assessing database coverage across different environments

We estimated the species-level coverage of the *MIDAS* database across metagenomes from different environments. Database coverage is defined as the percent (0 to 100%) of genomes from cellular organisms in a community that have a sequenced representative at the species level in the reference database. We estimated database coverage by (i) computing the total coverage across all species in the *MIDAS* database by mapping metagenomic reads to 15 universal single-copy genes and applying species-level mapping thresholds, (ii) computing the coverage across all microbial species, including those absent from the *MIDAS* reference database using the tool *MicrobeCensus* (Nayfach and Pollard 2015), and (iii) taking the ratio of these two quantities, multiplied by 100. We applied this approach to metagenomes from human body sites (Consortium 2012), human stool (Qin et al. 2012; Li et al. 2014; Obregon-Tito et al. 2015; Rampelli et al. 2015), baboon stool (Tung et al. 2015), mouse stool (Xiao et al. 2015), ocean water (Sunagawa et al. 2015), and soil from deserts, forests, grasslands, and tundra (Fierer et al. 2012). To identify possible taxonomic groups that harbored novel species in the human gut, we performed Spearman correlations between database coverage and the relative abundance of genera, estimated using mOTU (Sunagawa et al. 2013).

### Tracking transmission of strains between hosts

We used marker alleles to track transmission of strains between hosts. We defined a marker allele as an allele at a genomic site that was present in only a single individual, or in the case of the mother-infant dataset, a single mother-infant pair. For simplicity, we only considered bi-allelic genomic sites. An allele was determined to be present in a sample if it was supported by ≥3 reads and ≥10% of the total reads mapped at the genomic site. These parameters were chosen to minimize the effect of sequencing errors and filter out low frequency variants that could not be consistently detected between samples. Marker allele sharing was computed as the percent of marker alleles in mother strains that were also found in her infant. To minimize variation in marker allele sharing due to sampling, we excluded individuals with fewer than 10 identified marker alleles for a species. We applied this procedure to 66 species found in stool metagenomes from 98 Swedish mothers and their infants (Backhed et al. 2015) as well as 123 American individuals from the Human Microbiome Project (HMP) (Consortium 2012). We included the HMP samples in order to increase sample sizes and therefore improve the specificity of marker alleles identified in mothers and their infants. As a positive control to assess the sensitivity of our approach, we quantified marker allele sharing for each species between pairs of technical replicates from the HMP. As a negative control to assess specificity, we quantified marker allele sharing for each species between pairs of unrelated individuals from the HMP, which were not used to identify marker alleles. Based on these results, we defined a transmission event as >5% marker allele sharing between a pair of individuals.

### Analysis of globally distributed marine metagenomes

To assess the global population structure of marine bacteria, we analyzed shotgun metagenomes collected from the *Tara* Oceans expeditions that corresponded to prokaryotic size fractions. We utilized up to 100 million reads per metagenome and analyzed only one sequencing replicate per sample. We used *MIDAS* to quantify the relative abundance of the 5,952 reference species, and based on these results identified 30 species that occurred at >3x sequencing depth in the greatest number of metagenomes. The least prevalent species was found in 23% of metagenomes. Next, we used *MIDAS* to quantify the gene content of these species across metagenomic samples. Reads were mapped to the pangenome database, and reads with <94% alignment identity were discarded. Mapped reads were used to compute the coverage of genes clustered at 95% identity. Gene coverages were normalized by the coverage of 15 universal single copy genes to estimate gene copy numbers. We estimated gene presence-absence by thresholding the gene copy numbers, whereby any gene with a copy number <0.35 was considered to be absent.

To uncover population structure, we performed a principle component analysis of the gene presence-absence matrix for each species. To assess the relationship between gene content and geography, we first quantified the PCA distance and geographic distance between metagenomic samples for each species. PCA distances were computed using the Euclidian distance between samples based on the first two principle components. Geographic distances were computed using the Great circle distance with the R package geosphere (Hijmans 2016). Mantel tests were computed using the R package vegan (Oksanen et al. 2016) to correlate the PCA distances to the geographic distances. Up to one million permutations were performed to assess significance.

## Software availability

*MIDAS* is implemented in Python and is freely available, along with documentation at: https://github.com/snayfach/MIDAS. Source code is additionally included as Supplemental Material. Our reference database of bacterial species and associated genomic data resources are available at: http://lighthouse.ucsf.edu/MIDAS

## Acknowledgements

We thank Jonathan Eisen and Dongying Wu for helpful discussions about marker genes and phylogenetics and Joshua Ladau for discussions about microbial biogeography. This project was supported by funding from NSF grant #DMS-1069303, Gordon & Betty Moore Foundation grant #3300, the San Simeon Fund, and institutional funds from Gladstone Institutes.

